# On Copiotrophy and Temperature: Controls on Microbial Maximum Growth Rate Versus Translation Rate Optimization

**DOI:** 10.1101/2025.11.04.686613

**Authors:** JL Weissman, Alexandra Walling, Emily J. Zakem

**Affiliations:** Institute for Advanced Computational Science, Stony Brook University, Stony Brook, NY, USA; Department of Ecology & Evolution, Stony Brook University, Stony Brook, NY, USA; Biosphere Sciences and Engineering, Carnegie Institution for Science, Pasadena, CA, USA

## Abstract

Maximum growth rate is often used as a primary axis of functional variation in studies of microorganisms because it is key parameter in models of microbial growth, because it is closely conceptually related to the “copiotrophy-oligotrophy” axis used to organize microbial functional roles within ecosystems, and because emerging tools make it straightforward to estimate from genomic and metagenomic data. However, temperature, via its influence on reaction kinetics, may act as a confounder in studies that measure genomic signatures of growth optimization across environments by decoupling the cellular optimization for relatively fast growth from absolute growth rates. Observations suggest that growth optimization need not always indicate rapid growth. For example, strong temperature gradients are the norm across much of the world’s oceans, where slow-growing deep-ocean microbes show elevated signals of genomic growth optimization relative to the faster-growing communities at the surface. Looking across environments, we find a negative relationship between genomic growth optimization and optimal growth temperature, indicating the potential decoupling of genomic traits associated with copiotrophy from maximum growth rate, particularly when measured in the presence of a strong temperature gradient. Our results suggest that, as a result of temperature’s confounding effects, genomic signatures of growth optimization better predict the ecological roles and functional genomic content of microorganisms than do growth rates themselves. Finally, we suggest reframing copiotrophy as a set of traits that allow an organism to escape from a thermodynamic baseline maximum growth rate, rather than in relation to a specific rate cutoff.

## Introduction

A persistent problem for linking microbial process to global-scale biogeochemical cycling in both terrestrial and aquatic environments is that microbial communities are incredibly functionally and taxonomically diverse, often comprising many hundreds or thousands of species [1–3]. Genomic trait inference, wherein microbial phenotypes are estimated directly from genomic and metagenomic data, provides a path forward to systematically organize microbial functional diversity [4, 5]. For example, estimating maximum growth rate from genomes groups organisms from across the microbial tree of life into fast- and slow-growing clusters with distinct evolutionary patterns and genomic functional content [6, 7]. Genomically-inferred growth strategies have also recently been shown to align closely with the global-scale biogeography and functional roles of marine picoheterotrophs [8].

Maximum growth rate is often included as a genomically-tractable axis of functional variation in comparative studies [6, 9], because variation in maximum growth rate is conceptually related to the “copiotrophy-oligotrophy” axis used by many to organize microbial functional roles within ecosystems. Under this framework, fast-growing “copiotrophs” are optimized for high-resource environments and slow-growing “oligotrophs” are optimized for low-resource environments [7, 10]. Specialization for environments in which resources are abundant versus scarce need not necessarily align directly with microbial growth rate [6], because other parameters also matter for such fitness [11, 12], but fundamental rate-yield tradeoffs in ATP production pathways suggest that these two features of microbial lifestyle (resource abundance specialization and maximum growth rate) should often be correlated [13, 14], and indeed metabolic traits associated with canonical “copiotrophs” also tend to be associated with fast growth [6, 7]. Other trait-based frameworks for characterizing microbial diversity include additional axes like stress-response, but still emphasize a core tradeoff between growth rate (or “resource acquisition”) and efficiency of resource utilization that maps onto the copiotroph-oligotroph dichotomy [15, 16].

Genomic approaches allow for direct estimation of microbial growth potential and thus potentially facilitate placement of organisms along the copiotrophy-oligotrophy axis [6, 9]. Several genomic features thought to be associated with optimization for rapid translation have also been shown to be associated with rapid growth, including elevated codon usage bias of highly expressed genes, high rRNA and tRNA gene copy numbers, increased translation initiation start sites, and the presence of elongation factor P and a reduction in polyproline sequences (Fig. S1a-c; [9, 14, 17, 18]). Each of these features is associated with a specific mechanistic hypothesis (*e.g.,* that codon optimization allows for rapid translation which allows for faster biomass production, that polyproline sequences stall translation and thus slow down this process without the assistance of elongation factor P). More broadly, translation optimization can be thought of as an indicator of high ribosomal investment over evolutionary time, which should correlate with growth rate according to fundamental laws of bacterial growth [19, 20]. These features, most commonly codon usage bias and rRNA copy number, are frequently used in statistical models that predict the maximum growth rate of an organism from its genome sequence [6, 9, 14].

This simple picture is complicated by the fact that factors unrelated to resource abundance can exert substantial control on absolute maximum growth rates, including temperature [9, 21] and substrate quality [8], in ways that are orthogonal to the classical copiotrophy-oligotroph spectrum of substrate availability [8]. For example, temperature strongly constrains growth through basic thermodynamics; reactions proceed faster at higher temperatures, barring enzyme denaturation. Thus, temperature constrains microbial growth such that organisms with clear patterns of genomic optimization for rapid growth in relatively colder environments will in reality grow much more slowly than organisms showing no such patterns of optimization that live in warmer environments. Looking across sequenced microbial genomes, accurate estimates of an organism’s maximum growth rate on the basis of genomic data require an estimate of optimal growth temperature [6, 9]. Similarly, if the metabolism of an organism produces relatively little ATP (*e.g.,* methanogenesis), even a great deal of translation optimization may not increase growth rates to the levels seen in organisms with more energetically favorable metabolic pathways. In other words, copiotrophy need not always mean fast growth. One example of this decoupling comes from considering a gradient in the “quality” of organic substrates fueling heterotrophic microbial growth, where in marine systems it has been shown that organisms living deep in the water column, believed to be consuming abundant but recalcitrant (low quality) carbon sources, have enhanced genomic signals of translation optimization even though their maximum growth rates are necessarily very low [8]. This slow growth reflects the less-favorable energetics of the metabolisms of these organisms as well as the fact that these organisms live in cold environments [8, 22]. Temperature in particular is a potential confounder across diverse study systems, especially given the increase in research interest studying organismal responses to a warming climate.

Here, we address how temperature gradients can confound inferences made about the evolution of a microbial population’s life history strategy, *i.e.,* copiotrophy, from maximum growth rates. We demonstrate that over gradients in temperature, genomic measurements of growth potential, such as the degree of codon usage bias across an organism’s highly expressed genes [9, 22], may, in fact, tell us more about the ecology of an organism than the maximum growth rate itself. In particular, we demonstrate that potentially misleading interpretations of maximum growth rate may be gleaned when there exists a negative correlation between organismal optimal growth temperatures and genomic signatures of translation rate optimization. We additionally show that such a relationship between these two traits is the norm rather than an exception. In such situations (e.g., along temperature gradients), we suggest that ecological patterns are better recovered by genomic indices of growth optimization, rather than direct growth rate measurements, whether maximal or instantaneous, in line with recent work showing that these two pieces of information about an organism do not always align in the environment [23].

## Results & Discussion

### A Simple Model of Growth Optimization as Divergence from a Thermodynamic Baseline

Maximum growth rates are expected to increase with optimal growth temperature (*T_opt_*) [9, 24–26], and this pattern is also represented in commonly used statistical models for maximum growth rate prediction. Together, genomic maximum growth rate prediction tools [6, 9] generally use the following, or very similar (*e.g.,* with a Box-Cox transformation in place of a log transformation), model for maximum growth rate (𝜇*_max_*):

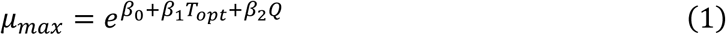

where Q is some measure of translation rate optimization (typically codon usage bias or ribosomal gene copy number) and the coefficients 𝛽_𝑖_ are fit using data drawn from the literature or experiments. Alternatively, we introduce a minor modification that turns this model into the Arrhenius curve:

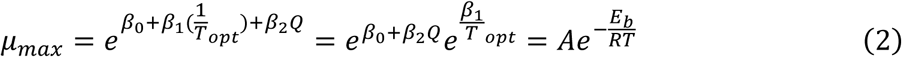

where 𝐴 = 𝑒^𝛽_0_+𝛽_2_𝑄^ is the pre-exponential factor and 𝛽_1_ = −𝐸_𝑏_/𝑅. This alternative model fits experimental growth and temperature data slightly better than Eq. 1 (gRodon [6] training data, see Methods; linear regression with a log transform, Eq. 1 AIC=1369 vs. Eq. 2 AIC=1333), suggesting that microbial growth kinetics across species scale with temperature in a similar manner to reaction kinetics, but also that growth kinetics can be modified by adaptations allowing rapid translation beyond this baseline expectation (where *Q* is greater than zero; see S1 Text for further model analysis and discussion). In Fig. 1a we see this expectation represented as a dashed line with a negative slope where codon usage bias is zero in our fitted model, such that organisms may have predicted maximum growth rates above this line if their associated value for scaled codon usage bias is above zero. In other words, the relationship with *T_opt_* sets the baseline for the maximum growth rate, and optimization for faster growth (*e.g.,* via genomic traits increasing the rate of translation) is an independent factor that raises 𝜇_*max*_ above this baseline. This result contrasts with within-organism growth kinetics, where the Arrhenius curve does not fit well across the full range of growth temperatures a single organism may encounter [24, 27], since growth rates ultimately decline beyond a given species’ *T_opt_*. It is known that an Arrhenius (*i.e.,* exponential) model better captures between-species patterns in temperature optima because each species is adapted for that particular optimum, thus mitigating any impact of protein denaturation or other negative fitness effects that may occur as temperatures increase [21, 28]. See S1 Text for further discussion of this model.

**Figure 1:**
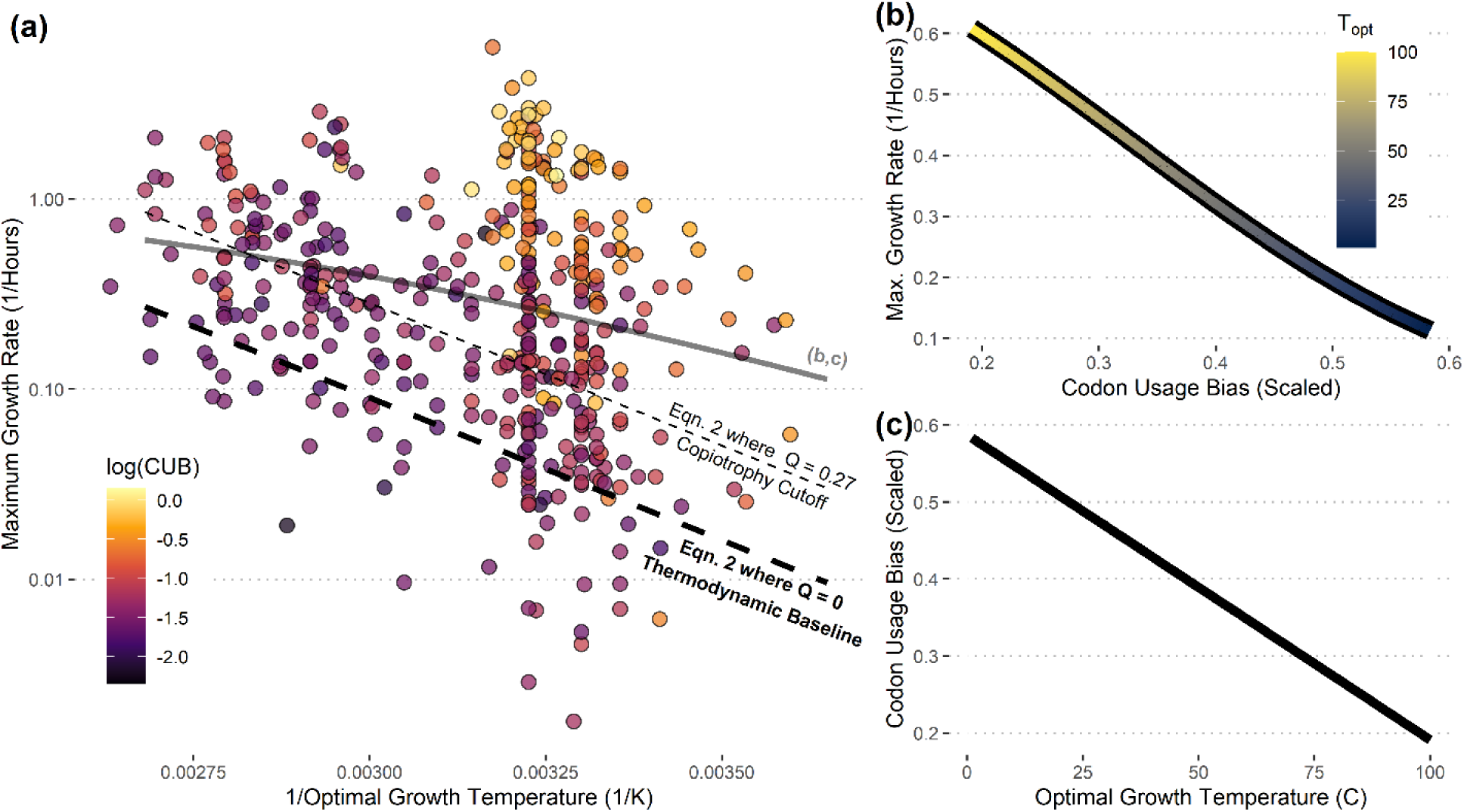
The growth rate versus translation optimization relationship can reverse along temperature gradients where translation optimization and optimal growth temperature are anti-correlated. (a) Maximum growth rate and T_opt_ data for species taken from the literature with scaled codon usage bias of the ribosomal proteins for corresponding genomes overlaid. Dashed lines show the fit of the modified Arrhenius model from Eq. 2 for the theoretical maximum growth rate versus 1/ T_opt_ relationship for organisms without any translation optimization (scaled CUB of zero, i.e., Q=0) as well as for organisms at the hypothetical oligotrophy/copiotrophy cutoff (based on clustering of species in Weissman et al. [6]). (b,c) Following Eq. 2, if we sample a series of organisms along a range of T_opt_ for which codon usage bias (Q) and T_opt_ are inversely related, it is possible that codon usage bias and maximum growth rate will also be negatively related across these organisms. The line shown in (b,c) corresponds to the solid line in panel (a).

Consider the hypothetical scenario where we sample organisms along a temperature gradient where the *T_opt_* of an organism is inversely related to its codon usage bias. This is potentially a good approximation of microbial populations in the surface ocean, where lower-latitude microbes are adapted to low nutrient supply in warmer waters and high-latitude microbes are adapted to high nutrient supply in colder waters (indeed, in the oceans, productivity is often correlated with cooler water temperatures because it relies on the upwelling of cold, nutrient-rich waters from depth.) In this case, one will likely obtain a series of organisms for which maximum growth rate appears to be negatively correlated with codon usage bias (Fig 1b,c), despite a positive mechanistic link between these two values encoded in our growth model. We show one example of this in Fig. 1, but this behavior is theoretically observable over a wide range of possible negative relationships between *T_opt_* and *Q* (S1 Text). In short, when sampling along a temperature gradient it is possible to reverse the observed relationship between translation optimization and growth potential, even if the overall mechanistic relationship does not change, if *Q* and *T_opt_* are negatively correlated across organisms.

### Translation Optimization is Frequently Negatively Correlated with Optimal Growth Temperature

We analyzed a database of 12,553 genome-matched species’ optimal growth temperatures drawn from the literature, as well as 112,441 genomic predictions of optimal growth temperatures for species representative genomes from GTDB v220 [29–31]. We found that genomic signatures of translation optimization were less likely to appear at hotter optimal growth temperatures than colder growth temperatures, with psychrophilic and mesophilic organisms showing high levels of translation optimization and thermophiles showing a striking absence of any such signals (Figs. 2a-c and S2). This negative relationship was robust to phylogeny (phylogenetic linear regression; codon usage bias, p=2.4e-47; rRNA, p=0.02; EFP, p 2e-3; Fig. S3) and was apparent across a wide variety of habitat types (Fig. S4), although was notably absent in host-associated organisms for which temperature ranges tended to be narrower (Fig. S5). Nevertheless, the negative translation optimization versus temperature relationship persists even among organisms that live in habitats that never reach high temperatures (*e.g.,* those above approximately 40C; Fig S4). This negative relationship was strongest among fast-growing organisms (Fig. S2), as there were many slow-growing organisms with little translation optimization present across the range of optimal growth temperatures (Fig 2a-c).

**Figure 2:**
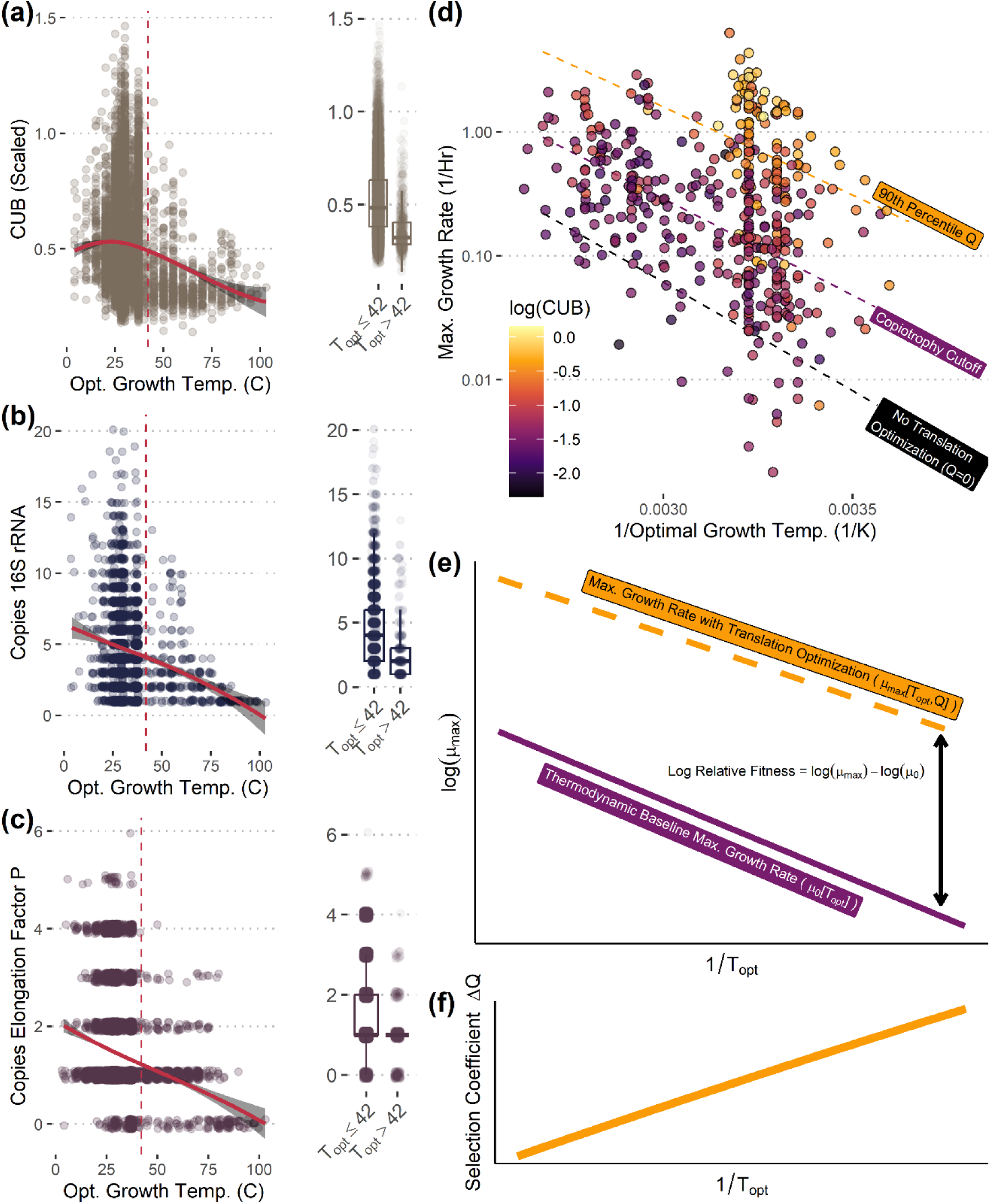
Translation optimization is negatively correlated with optimal growth temperature across species. (a-c) Multiple genomic signatures associated with optimization for rapid translation are negatively correlated with an organism’s T_opt_. Red lines depict a GAM fit. Temperature data from a large species-level database of genome-matched T_opt_ from the literature (n = 12,553). (d) The fit of a diminishing returns model of translation optimization overlaid on data as in Fig .1a, with dashed lines depicting model fit for different values of codon usage bias (no bias, the theoretical threshold between copiotrophy and oligotrophy as proposed in [6], and the 90^th^ percentile of codon usage bias based on the data shown in this panel). (e-f) Conceptual figures showing how relative fitness is defined under the diminishing returns model and how the selection coefficient for an increase in translation optimization changes with increasing T_opt_ under this model.

The cause of this general negative relationship is not immediately apparent. According to our base Arrhenius model, the relative fitness benefit of increasing translation optimization is constant across temperatures (S1 Text). We considered three potential hypotheses for why organisms with higher optimal growth temperatures might evolve weaker genomic signals of translation optimization. First, we hypothesize that lower effective population sizes in heat-loving organisms may lead to less efficient selection. Genomic signatures like codon usage bias are the result of many mutations with small fitness effect, so that if *s<<1/Ne*, where *s* is the coefficient of selection for one of these small effect mutations and *Ne* is the effective population size, drift might overwhelm the evolution of translation optimization. While *Ne* is difficult to estimate in microbial populations [32], habitats that reach temperatures greater than 40C are less common and frequently are produced by geological (*e.g.,* hot springs) or metabolic (*e.g.,* compost heaps) sources of thermal energy rather than solar- or host-derived heat (Fig. S6). Nevertheless, low effective population size alone cannot explain why the negative correlation between *T_opt_* and translation optimization persists among organisms with *T_opt_* below 40C or that live in habitats that never reach that temperature (Fig S4).

Second, we hypothesize that constraints on growth rates unrelated to translation optimization could lead to diminishing returns for increased translation optimization as the maximum growth rate of an organism increased, and that this could in turn lead to decreased selection for translation optimization at higher growth temperatures. In other words, the degree of translation optimization may have a smaller impact on maximum growth rates at higher temperatures than lower temperatures. Such a pattern could occur if some other process involved with growth became rate-limiting as translation rates and/or ribosomal investment increase. We incorporated a diminishing return directly into our base Arrhenius model (Fig 2d-e; S1 Text), which confirmed that the selection coefficient for increased translation optimization would be inversely related to *T_opt_* under a diminishing return scenario (Fig 2f; S1 Text).

Finally, it is also possible that faster growth itself could simply be less beneficial at higher temperatures, decreasing the selection coefficient for increases in *Q*. In other words, rather than diminishing returns, there may be no evolutionary incentive to optimize for faster growth at these temperatures for other reasons. For example, if thermophiles as a rule experienced a stronger tradeoff between carbon use efficiency and growth rate than mesophiles, this could lead to weaker signals of translation optimization in thermophiles. Some ecological models also suggest that slower growth should be favored at warmer temperatures in line with this hypothesis [33, 34].

In reality, it is likely that some combination of these three hypotheses leads to the observed negative correlations between *T_opt_* and *Q* (Fig 2a-c), though insufficient data makes it challenging to distinguish them at this time.

### Translation Optimization and Thermophily Rarely Co-Occur in an Organism

Above a temperature threshold of 42C, we observed few organisms with signs of translation optimization (Figs. 2-3 and S2). Corkrey et al. [25] noted an absence of fast-growing organisms with *T_opt_* in the range 42C-60C, which they call the “thermophile-mesophile gap” (Fig 3a). Viewed alongside information about organismal codon usage bias and our growth model, we propose that this gap exists between two kinds of fast-growing organisms. Below 42C, organisms with fast maximum growth rates are those with high codon usage bias (Fig 3a,b). Above 42C, organisms do not have high codon usage bias, but as optimal growth temperatures increase beyond 60C basic thermodynamics leads to faster growth. The “gap” between these two classes of fast growers is where high optimal growth temperature cannot compensate for low translation optimization. In other words, there are two ways an organism may increase its maximum growth rate – either by increasing its optimal growth temperature or by optimizing its genome for rapid growth (*e.g.,* by increasing the rate of translation of highly expressed genes), but organisms rarely if ever seem to combine these two traits (Fig 3b,c).

**Figure 3:**
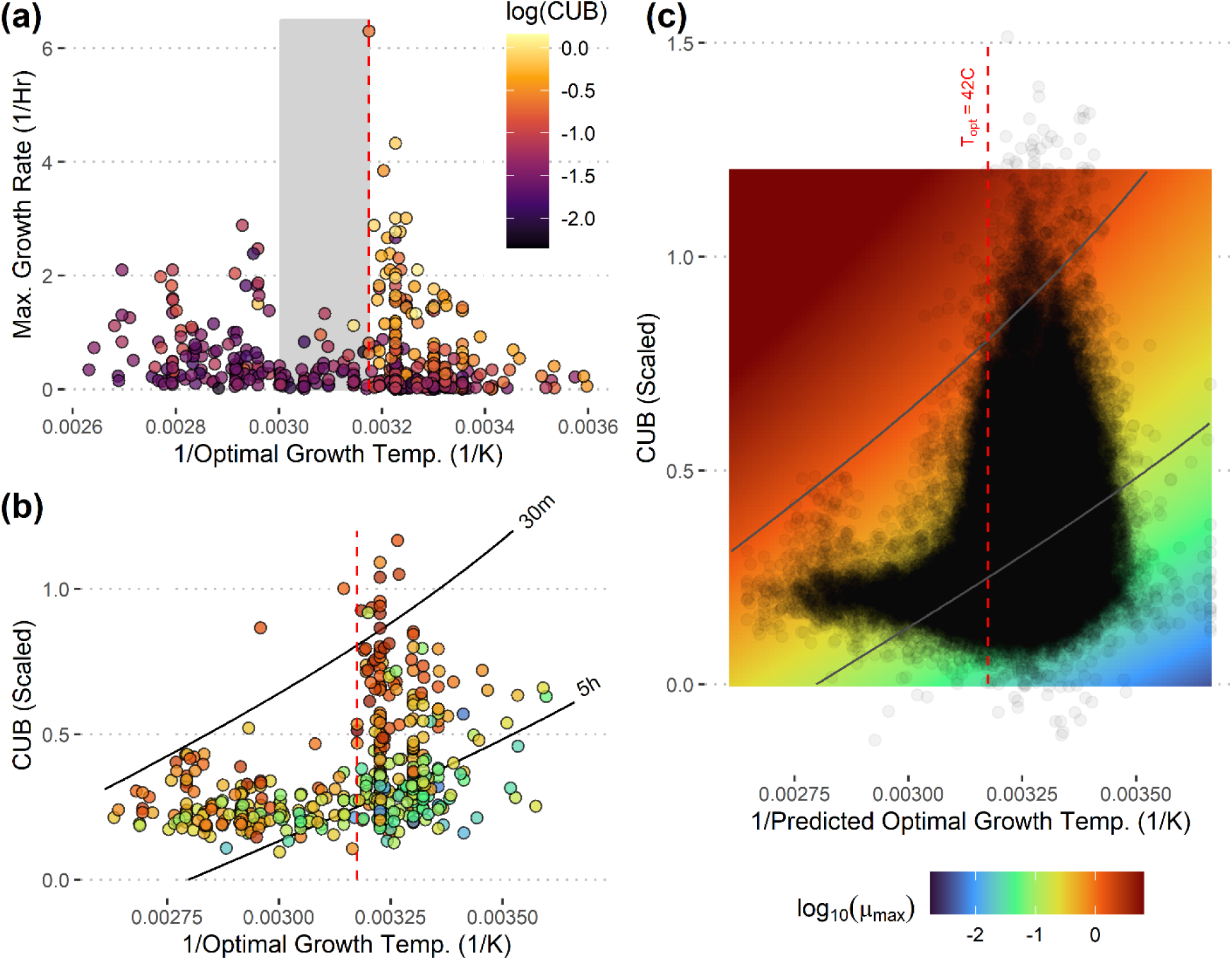
Fast-growing organisms either have high translation optimization or a high optimal growth temperature, but very rarely both. (a) Data as in Fig. 1a, plotted on a linear y-axis. Gray shaded region denotes the hypothetical thermophile-mesophile gap between 42C and 60C where fast-growing organisms are absent from this and other datasets [21–22]. Vertical dashed red line at 42C. (b) The same dataset plotted on different axes, shows that organisms with T_opt_>42C almost never have elevated codon usage bias. Solid lines show contours from the translation optimization diminishing returns growth model (S1 Text) at a minimum doubling time of 5 hours and 30 minutes. Notice that organisms near the 30 minute contour either have high T_opt_ or a high codon usage bias, but not both. Color fill shows maximum growth rates from the literature (experimental). (c) Species representative genomes from GTDB v220 mapped onto the space in (b) (n=112,441), with predicted maximum growth rates for each combination of codon usage bias and T_opt_ from the diminishing returns model shown as a color gradient in the background (same scale for panels b,c; empricial rates in b and predicted rates in c).

### Translation Optimization for Understanding Ecological Temperature Gradients

Given the above results that (1) negative *T_opt_* versus translation optimization relationships can cause misleading negative correlations between translation optimization and maximum growth rates (Fig 1) and (2) that such negative relationships are common (Fig 2a-c, S3-S4), we sought to find examples of these patterns in environmental datasets. For example, looking across depths in the global oceans using metagenomes from the BIOGEOTRACES dataset and temperature-corrected community-level average maximum growth rate predictions from gRodon [35], we found that codon usage bias and maximum growth rate are anti-correlated across depths, due to a strong temperature gradient (Fig. 4a-c). In reality, two factors work to limit the growth rates of organisms living deeper in the water column: temperature, which we discuss here, and nutrient quality, which specifically limits the growth rates of marine heterotrophs at depth [8]. Moving beyond marine systems, we also examined a dataset of *streptomyces* sister taxa sampled from a latitudinal gradient of soils across North America [36], and saw a similar pattern where isolates from a Southern phylogroup had warmer *T_opt_* and decreased codon usage bias relative to a Northern phylogroup, leading to a negative correlation between codon usage bias and maximum growth rate across these groups (Fig S7).

**Figure 4:**
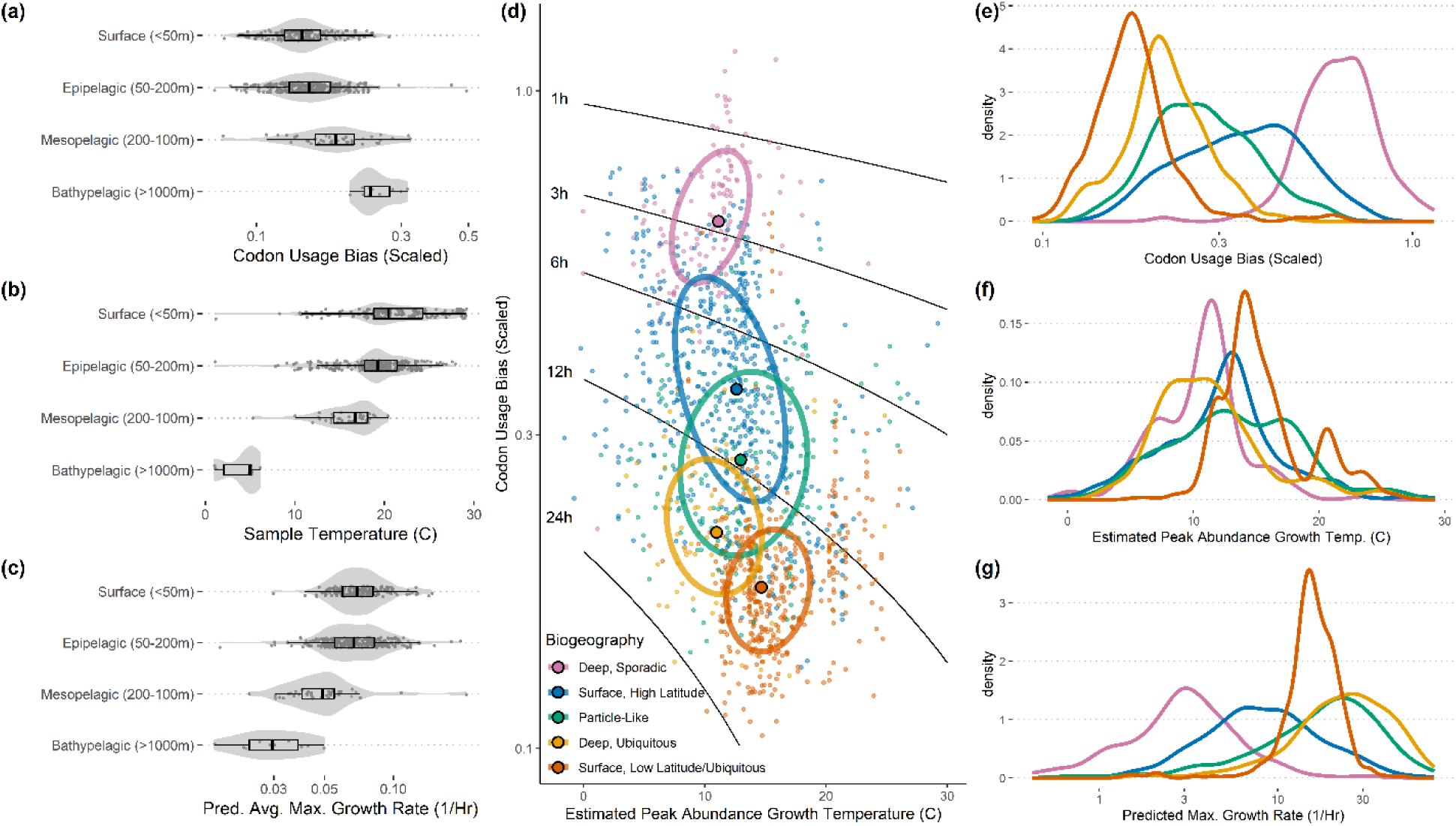
Translation optimization, not maximum growth rate, predicts the distribution of ecological strategies in the presence of a strong temperature gradient. (a-c) Average community-wide codon usage bias, sample temperatures, and predicted average maximum growth rates from BIOGEOTRACES metagenomes. Note that codon usage bias and predicted maximum growth rate are anti-correlated across depths. (d-f) Biogeography for marine picoheterotrophs appears to organize along a gradient of translation optimization (CUB) rather than growth rate or temperature. Data taken from [8] where genomes from GTDB v220 were mapped to biogeographic categories along a transect of the Pacific Ocean running from 60N to 60S with depth profiles up to 600m using high-resolution 16S rRNA profiles. (d) Each point is a genome, with color representing the assigned biogeographic category, and ellipses are drawn at the 50^th^ percentile for each category with centroids shown as larger outlined points. Solid black lines show contours of the predicted min. doubling times associated with each estimated growth temperature and codon usage bias. (e-f) The marginal distributions of the categories in panel (d) for the three axes shown in that panel.

If maximum growth rate and translation optimization do not always covary within an environment, it is reasonable to ask whether one of these two traits is more useful than the other for understanding biogeography. Using data mapping GTDB v220 representative genomes to biogeographic classes across a Pacific Ocean transect (Fig 7; [8]) we found that biogeographic classes appeared to be primarily arranged along an axis of translation optimization rather than *T_opt_* or maximum growth rate (Fig. 4d-g). This suggests that by grouping microbes by their maximum growth rate in the presence of temperature gradients we may lose information about the forces structuring ecosystems.

### Metabolic Traits Primarily Co-Vary with Translation Rate Optimization

Given the two paths organisms may take to achieve rapid growth (Fig. 3), we predicted that functional genes related to copiotrophy (*e.g.,* those facilitating rapid acquisition of resources) would be most beneficial to organisms that achieve fast growth through translation optimization and would be proportionally less beneficial for organisms already capable of rapid growth by nature of life in a warm environment. That is, the benefit of carrying genes allowing for the rapid exploitation of resources would be seen most dramatically in growth-rate maximizers who are also translation-rate optimizers but would be minimal for growth-rate maximizers who simply live in warmer environments.

We built linear models of gene family presence/absence for family-level representative genomes from GTDB v220 using codon usage bias and *T_opt_* as predictors (Fig 5a, S9). Across models showing significant interactions (p<0.01 for both coefficients after Benjamini-Hochberg correction), we found that most gene families (56%) were positively associated with codon usage bias and negatively associated with *T_opt_*, corresponding to growth-maximizers who are also translation-optimizers but not temperature-optimizers (Fig 5a, S9). In contrast, only a small minority of gene families (16%) were found to be positively associated with growth in general (positive coefficients for both codon usage bias and *T_opt_*). While in most cases gene families with significant coefficients were of unknown or only generally annotated functions, gene families that had a positive codon usage bias relationship and a negative *T_opt_* relationship were enriched for functions involved in translation, amino acid transport and metabolism, and ion transport and metabolism, among others (Fig. 5a). We repeated the same analysis at the level of metabolic pathways, and found that the presence of metabolic pathways often conceptually associated with copiotrophy such as carbohydrate and amino acid metabolism were positively associated with codon usage bias and negatively associated with *T_opt_*, whereas canonically oligotrophic pathways like methanogenesis and hydrogen oxidation are negatively associated with codon usage bias and positively associated with *T_opt_* (Figs. 5b-e and S8). Together, these results suggest that copiotrophic traits are correlated with growth optimization, but not necessarily maximum growth rate itself due to confounding temperature effects.

**Figure 5:**
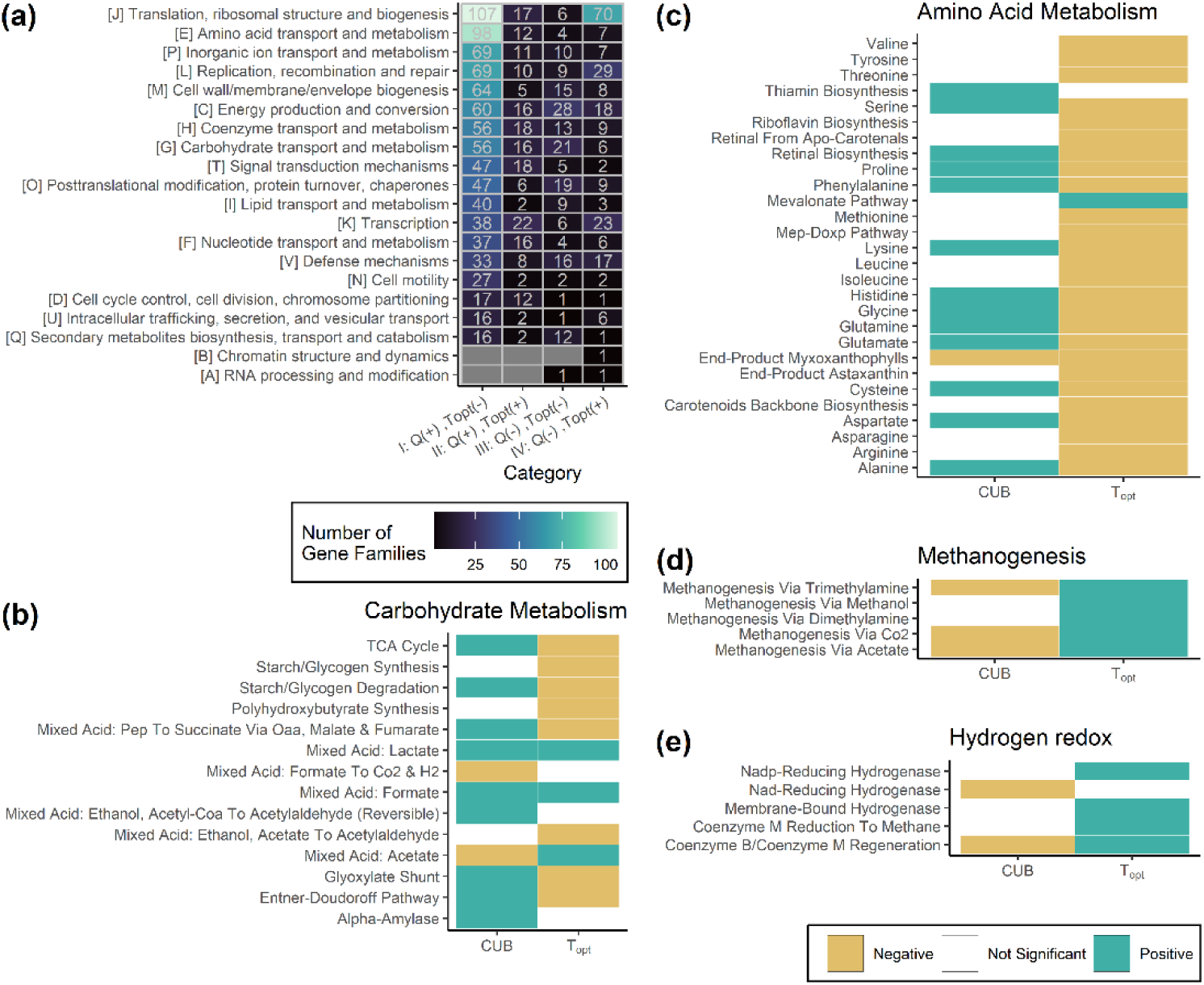
Genomic translation optimization predicts the presence of metabolic pathways associated with copiotrophy. (a) The functional breakdown of CUB and OGT associated gene families. Labels on x-axis indicate whether there was a positive or negative association with translation optimization (Q) and optimal growth temperature (T_opt_), corresponding to the categories in S9 Fig. Notably, the large majority of genes have unknown function or only a general functional prediction, whereas genes families that had positive relationships with translation optimization and negative relationships with temperature (i.e., being primarily translation associated rather than being correlated with max. growth rate) are more likely to be involved in translation, transport, and biogenesis. (b-e) At the pathway level, canonically copiotrophic pathways like carbohydrate and amino acid metabolism tend to be positively associated with codon usage bias, whereas canonically oligotrophic metabolisms like methanogenesis and hydrogen redox are positively associated with growth temperature (p<0.01, Benjamini-Hochberg correction).

## Conclusions

We found that negative relationships between genomic translation optimization and optimal growth temperature are widespread across taxa and environments (Figs. 2-3,S2-S4), and showed how such relationships can and do lead to a decoupling of genomic traits that enable rapid growth and actual observed maximum growth rates in the presence of strong temperature gradients (Figs. 1,4,S7). As a result, genomic translation optimization is a better predictor of canonically “copiotrophic” metabolic traits than maximum growth rate itself looking across sequenced genomes (Figs. 5, S8, S9).

In some environments, it is possible that the inversion of the translation-optimization versus maximum growth relationship leads genomic traits we expect to be associated with “copiotrophy” to become anti-correlated with growth. Considered out of context, this could lead one to propose certain functional relationships between, *e.g.,* gene abundances and growth that were entirely spurious. Similarly, using genomic translation-optimization (*e.g.,* number of 16S rRNA copies, codon usage bias) to infer maximum growth rate across a gradient like depth or latitude in marine systems without a temperature correction could lead to a reversal of the true maximum growth rate relationship in one’s inferences. Importantly, this confounding effect is as real for growth rates measured directly in the field or lab as it is for growth rates predicted from genomes. In both cases, temperature can act as a major confounder when trying to link growth to gene content or ecological role. These arguments generalize to instantaneous rates (*e.g.,* measured by qSIP [37]) which are limited by temperature. The most detailed measurements may not tell us much about the composition of ecological strategies in a community if not discussed in the context of local temperature.

Following this line of argument, we suggest reframing copiotrophy as a set of traits that allow an organism to escape from a thermodynamic baseline maximum growth rate. That is, we can think of degrees of copiotrophy as how far away an organism’s maximum growth rate is from its expected growth rate at the same optimal growth temperature in the absence of translation optimization (Fig 1a, Q=0 line). Thermophiles, though they may grow fast, generally would not be considered copiotrophs because the baseline expectation for growth is already so high. Alternatively, even slow-growing psychrophiles might show genomic signatures associated with copiotrophy. Conveniently, measures of translation optimization appear to directly measure this “off-baseline”, suggesting that they are natural measures of copiotrophy. Our previous work showed, at least among marine heterotrophs, carbon lability sets an additional growth baseline from which organisms similarly diverge, as indicated by elevated codon usage bias in highly expressed genes, to take advantage of resource abundance [8]. Together, these results emphasize the importance of interpreting genomic signals of translation optimization like codon usage bias in the context of resource abundance, while other factors like resource quality and temperature may limit growth maxima in entirely orthogonal ways.

It is an open question as to why optimal growth temperatures are negatively correlated with translation optimization across organisms. We suggested three hypotheses: (1) that *Ne* and optimal growth temperature may be negatively correlated due to a scarcity of habitats with temperatures exceeding 42C (Fig S6), leading to less efficient selection in thermophiles, (2) that the benefit of mutations increasing translation optimization diminishes with increasing optimal growth temperature because the phenotypic effect of these mutations is smaller at faster growth rates (diminishing returns model, S1 Text, Fig. 2), or (3) that fast growth may be less beneficial among organisms with high optimal growth temperatures because of other tradeoffs with growth rate that could become stronger with temperature. Possibly the observed effect could be a product of all three hypotheses, though we believe that hypothesis two would be easiest to test directly in the laboratory with controlled competitions between mutant strains across a range of temperatures.

Finally, we note that the arguments we make about rate variation across a temperature gradient, while simple, are also generalizable to any temperature-dependent rate. That is, one might find the highest copy number of a gene that produces a given metabolite in the genome of an organism that lives at the coldest temperatures, rather than in the genome of an organism that produces that metabolite at the fastest rate. Similarly, two organisms may be equally reliant on the production of a metabolite and produce it at the same rate, but if one lives in a colder environment, it may need a higher gene copy number to sustain that production. Large-scale comparative genomic analyses that ignore how temperatures constrain metabolic rates will consistently miss, or even reverse, important gene-function relationships.

## Methods

For all bioinformatics analyses, default parameters were used unless otherwise specified. All data and scripts used to generate analyses and figures available at https://github.com/jlw-ecoevo/copio_temp/. For model details and analysis see S1 Text.

### Genomic Trait Database Annotation

We obtained the 113,104 species-level representative genomes from the GTDB v220 release, which includes a combination of genomes sequenced from isolates, high-quality metagenome-assembled genomes (MAGs), and high-quality single-cell amplified genomes (SAGs)[38–41]. All of these genomes were annotated using prokka v1.14.6 (using ‘--kingdom Archaea’ for archaeal genomes) [42]. Carbohydrate-active enzymes (CAZymes) were annotated using dbCan v2.0.11 with annotations retained if confirmed by at least two methods [43]. Optimal growth temperatures were predicted using GenomeSPOT v1.0.1 [30]. Maximum growth rates and codon statistics were estimated using gRodon v2.4.0 for the 112,441 genomes with sufficient numbers of annotated ribosomal proteins for confident prediction ([6, 22]; using metagenome mode to control for any potential contamination in the MAGs and SAGs and using temperature predictions from GenomeSPOT; Figs. 3c,S3,S4b,S5b). Species-level 16S rRNA counts were obtained from rrnDB-5.9 and mapped to 4,357 GTDB v220 representative genomes by species name [44]. For 12,553 species representative genomes we were additionally able to match experimentally-measured optimal growth temperatures in the Gosha database which are drawn from the literature [31] (Figs. 2a-c,S4a,S5a). A subset of one randomly selected representative genome per family (5,463 genomes) was also annotated using eggnogmapper v [45] and resulting gene families were grouped into pathways using KEGG-Decoder and KEGGaNOG v1.1.17 [46] (Figs. 5, S8).

### Other Trait Data

We used the mapping of 1,417 GTDB v220 representative genomes to marine biogeographic categories described in Zakem et al. [8] (Fig. 4d-g).

Genome assemblies for the 20 northern and southern phylogroup genomes described by Choudoir et al. [36] were annotated with predicted maximum growth rates and optimal growth temperatures using gRodon v2.4.0 [6] and GenomeSPOT v1.0.1 [30].

We also re-analyzed the original 17,347 genomes matched to 444 species with experimentally measured maximum growth rates and optimal growth temperatures from the literature taken from the Madin et al. [47] trait database that was used to train the original gRodon software (Figs. 1a, 2d,3a-b,S1,S2).

BIOGEOTRACES metagenomes [48] were annotated with maximum growth rates as described in Weissman et al. [22] (Fig. 4a-c). Temperatures were taken from BIOGEOTRACES metadata [48].

Surface temperature data in Fig S6 was obtained from NASA MODIS [49, 50] and body temperature data from Moreira et al. [51].

### Statistical Analysis

We performed phylogenetically controlled linear regression analyses using a Brownian motion model in the phylolm v2.6.5 R package [52] and the phylogeny of representative species genomes provided with the GTDB v220 release. The robustness of phylogenetic regressions to specific species and phyla, and to sample size, were assessed using the sensiphy v0.8.5 R package [53] which repeatedly drops data from the dataset to assess sensitivity.

Gene and pathway analyses were performed by building per-gene or per-pathway logistic regression models (glm in R v4.3.3 base stats [54]) with codon usage bias and optimal growth temperature as predictors and correcting for multiple testing using a Benjamini-Hochberg correction.

The growth model in Eq. 2 was fit to the original gRodon training data of maximum growth rates and optimal growth temperatures from the Madin et al. [47] trait database using multiple regression (lm in R v4.3.3 base stats). The diminishing returns model (described in S1 Text) was fit to the data using the nls function in R v4.3.3 base stats.

## Acknowledgements

E.J.Z. was supported by National Science Foundation grant (2125142).

