## Supplemental Figures for "On Copiotrophy and Temperature: Controls on Microbial Maximum Growth Rate Versus Translation Rate Optimization"

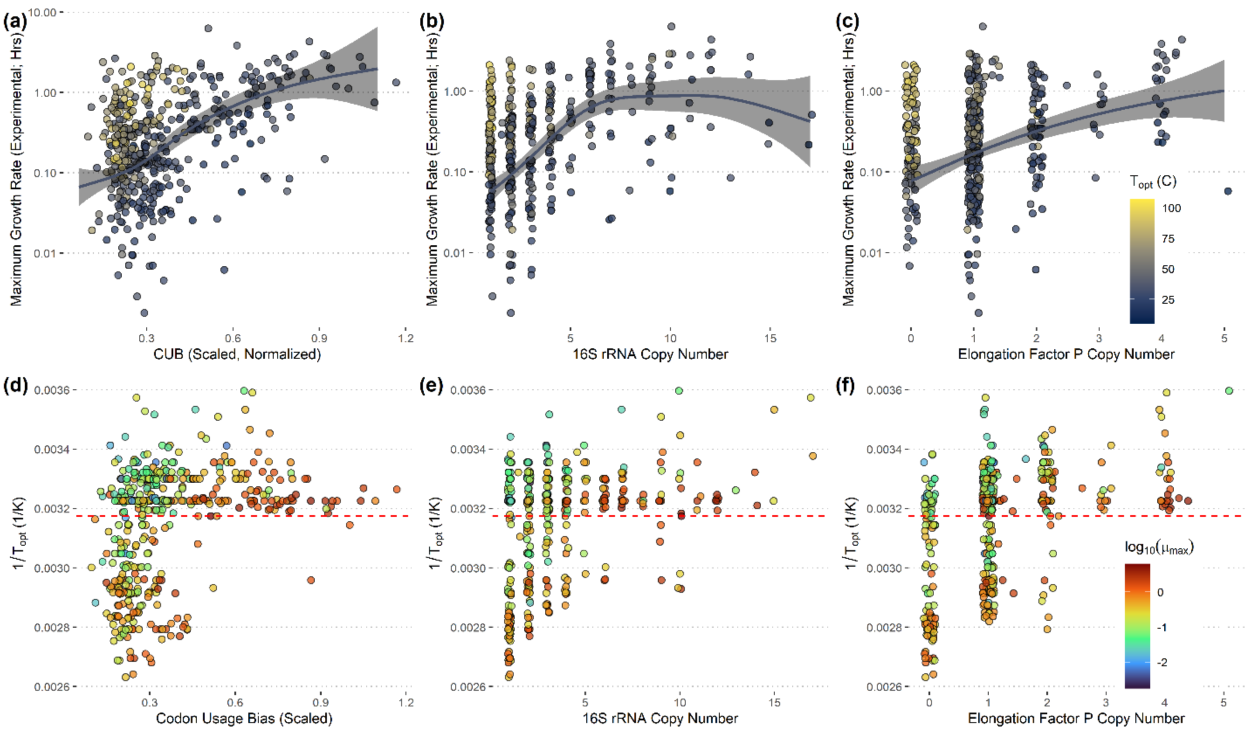


***Supplemental Figure 1:*** *(a-c) Maximum growth rate is positively associated with various measures of translation optimization, but organisms with high* $T_{opt}$ *defy this pattern. Solid lines depict GAM fit for organisms with* $T_{opt}$ *below 42C. (d-f) Organisms with high* $T_{opt}$ *rarely show signs of translation optimization on their genomes. Dashed red line at 42C.*

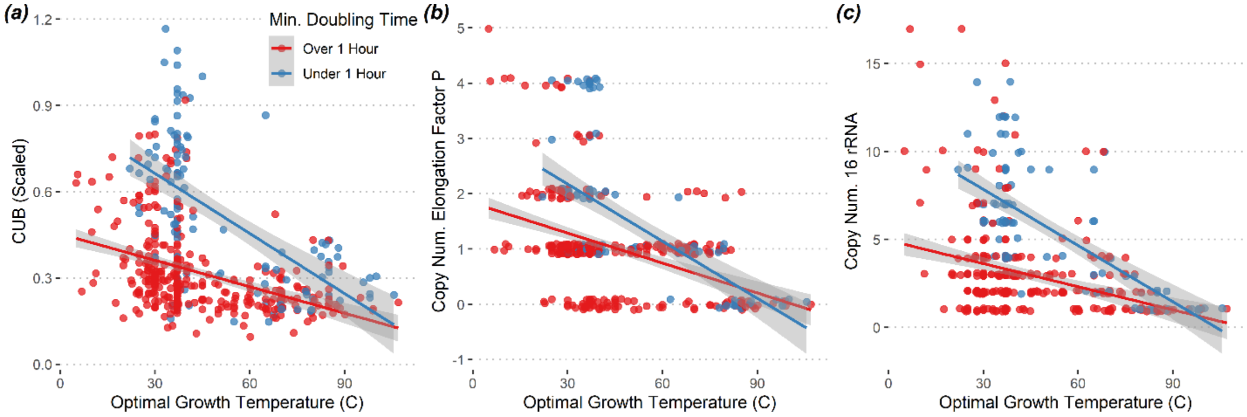


***Supplemental Figure 2:*** *The negative relationship between optimal growth temperature and translation optimization is strongest among fast-growing organisms.*

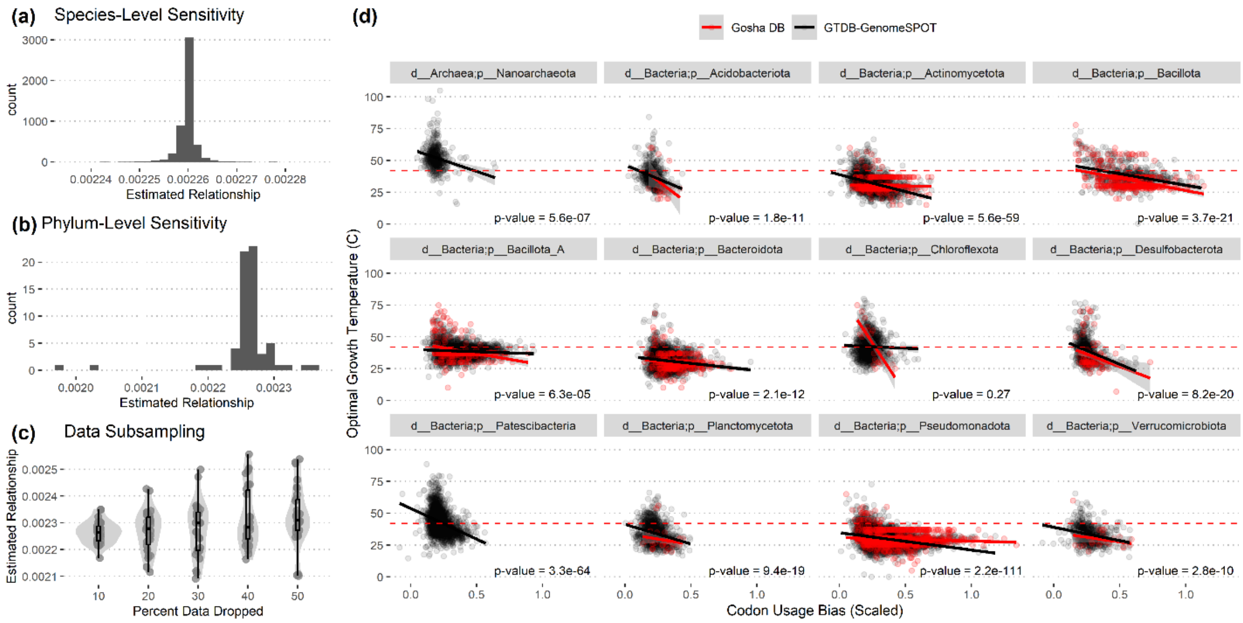


***Supplemental Figure 3:*** *The negative relationship between optimal growth temperature and codon usage bias is robust to phylogeny. (a) Phylogenetic linear regression sensitivity analysis to held-out species. (b) Phylogenetic linear regression sensitivity analysis to held-out phyla. (c) Phylogenetic linear regression sensitivity analysis to varying percentages of the dataset being held-out. (d) The relationship between codon usage bias and* $T_{opt}$ *for different phyla either using predicted* $T_{opt}$ *across all of GTDB v220 (black) or a database of growth temperatures from the literature (red). Phyla with at least 500 representatives in GTDB shown.*

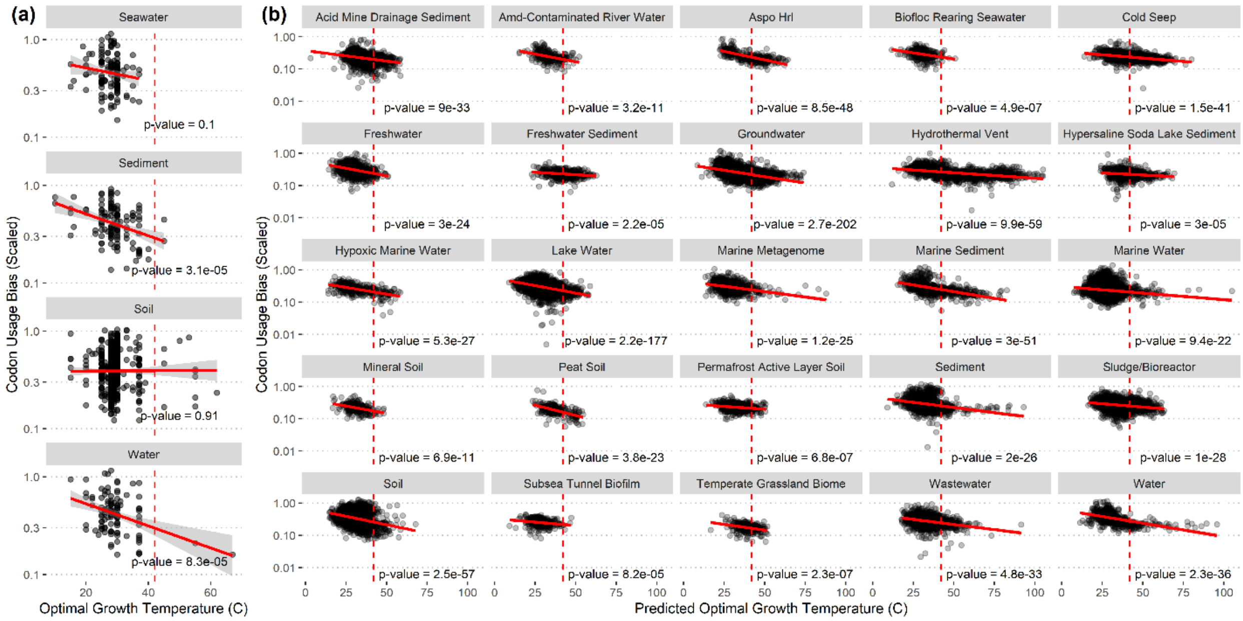


***Supplemental Figure 4:*** *The negative relationship between optimal growth temperature and codon usage bias is apparent across diverse environments. Data in panel (a) from a database of* $T_{opt}$ *measurements from the literature and in panel (habitats with >100 representatives shown) and in (b) from predicted* $T_{opt}$ *for GTDB v220 (habitats with >250 representatives shown). Habitat data taken from NCBI isolation source information, excluding host-associated habitats (see Fig S5).*

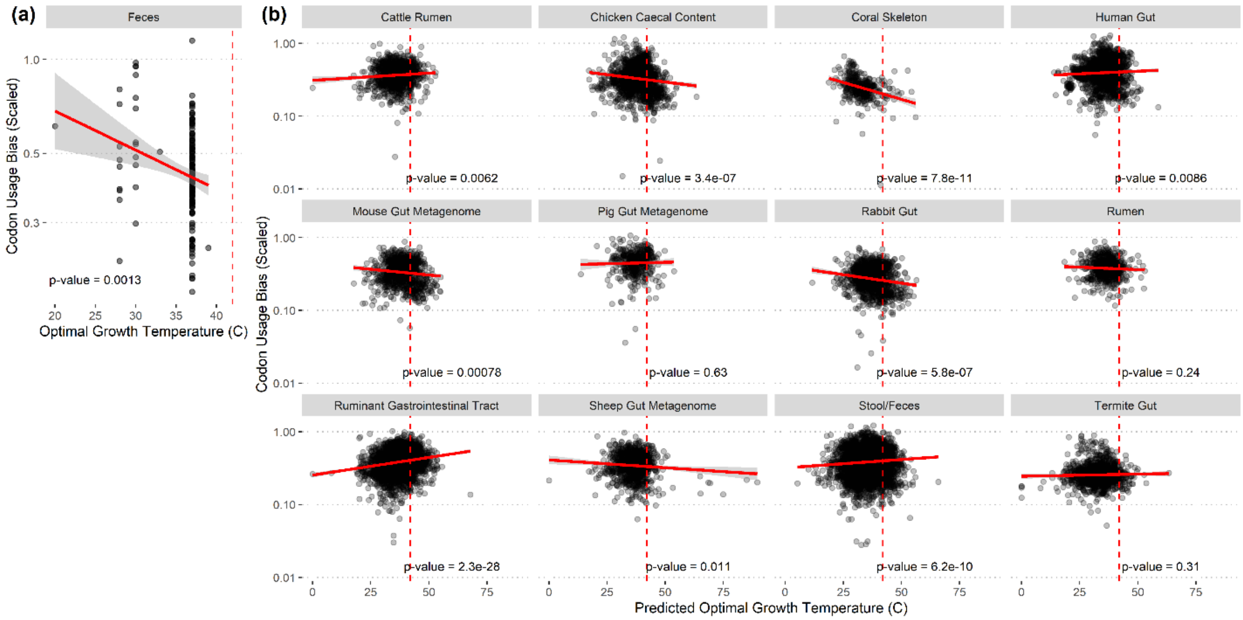


***Supplemental Figure 5:*** *The negative relationship between optimal growth temperature and codon usage bias does not consistently appear among host-associated environments. Data in panel (a) from a database of* $T_{opt}$ *measurements from the literature and in panel (habitats with >100 representatives shown) and in (b) from predicted* $T_{opt}$ *for GTDB v220 (habitats with >250 representatives shown)). Habitat data taken from NCBI isolation source information.*

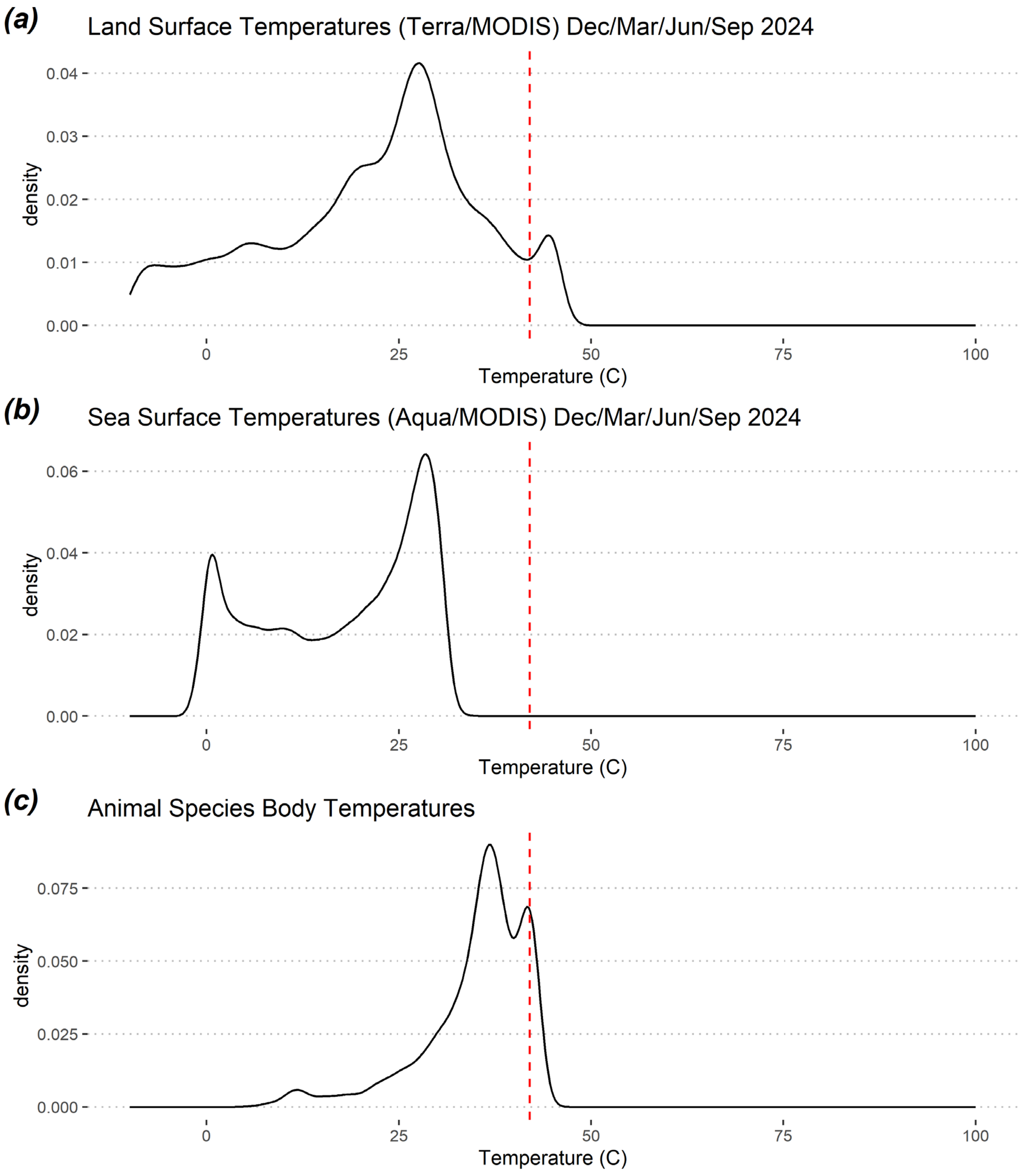


***Supplemental Figure 6:*** *Habitats with temperatures greater than 42C are rare. (a) Distribution of global land surface temperature measurements between -10C and 100C during four months of 2024 from NASA Terra/MODIS. (b) Distribution of global sea surface temperature measurements between -10C and 100C during four months of 2024 from NASA Aqua/MODIS. (c) Distribution of average animal body temperature measurements across species. Most high temperatures are associated with passerine birds.*

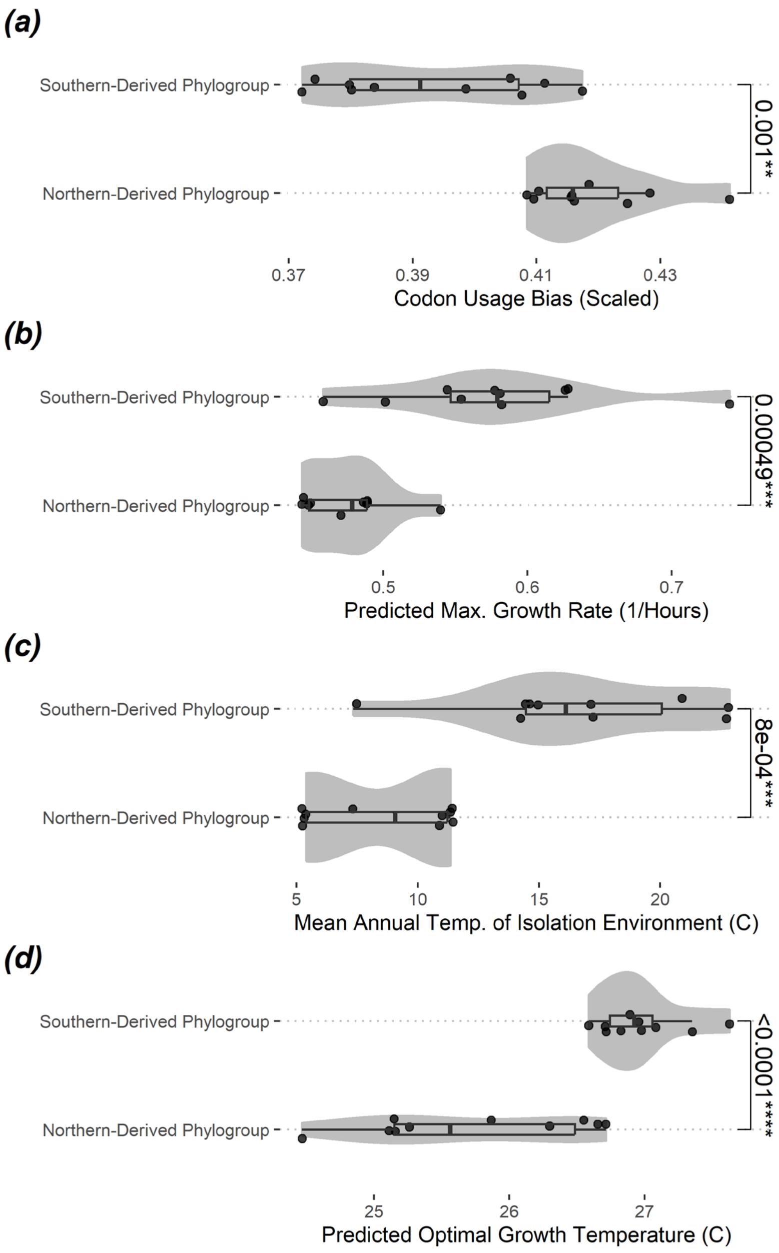


***Supplemental Figure 7:*** *Southern- and nothern-derived phylogroups of* streptomyces *collected from North America show anti-correlated codon usage bias and optimal growth temperature, as well as anti-correlated codon usage bias and predicted maximum growth rate. Significance levels shown from Welch's t-test. Optimal growth temperatures predicted with GenomeSPOT and growth rates predicted with gRodon.*

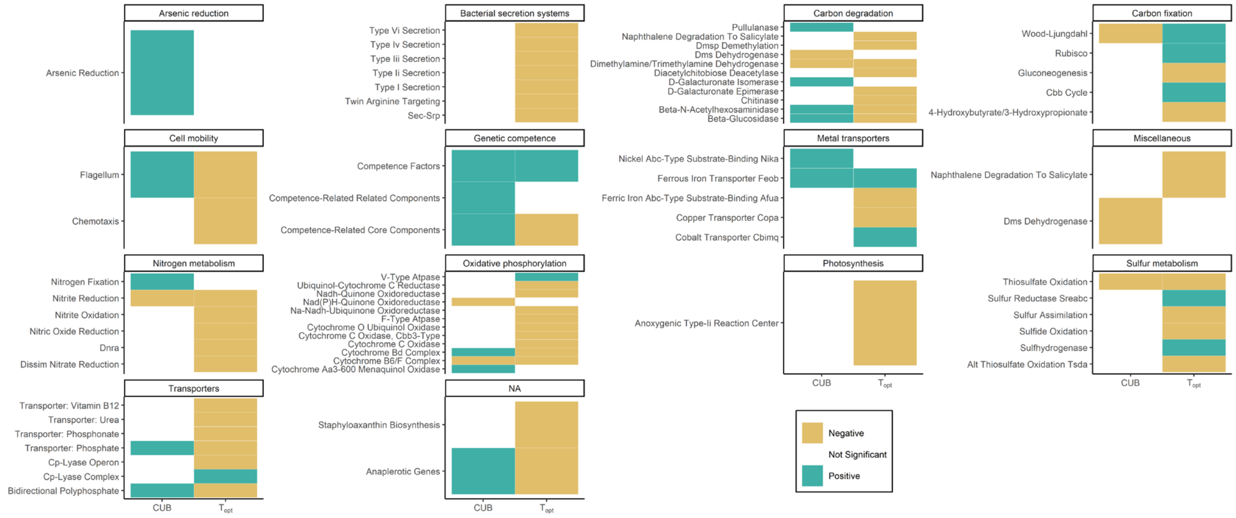


***Supplemental Figure 8:*** *Pathways associated with codon usage bias and/or optimal growth temperatures not shown in Fig 5.*

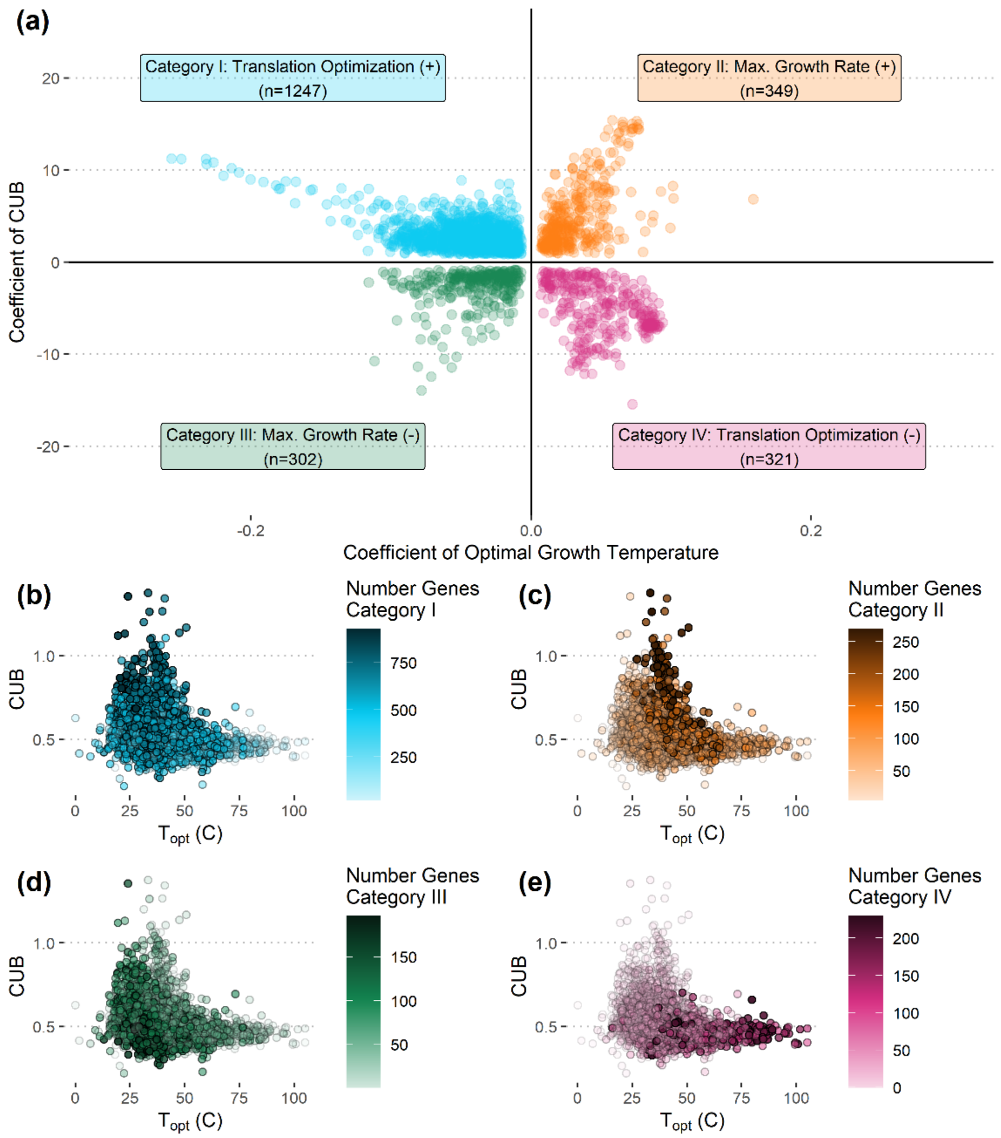


***Supplemental Figure 9:*** *(a) More gene families correlate with translation optimization (CUB) than maximum growth rate itself across families in GTDB v220. The coefficients of regression models of gene family (COG) presence/absence across genomes where codon usage bias and* $T_{opt}$ *are significant predictors (p<0.01, Benjamini-Hochberg correction) show that most gene families are positively associated with codon usage bias and negatively associated with* $T_{opt}$ *(category I), whereas only about 16% of the gene family models with significant relationships show a positive association with growth writ-large (category III; i.e., a positive association with both codon usage bias and growth temperature). (b-e) The distribution of CUB and OGT associated gene families across GTDB representative genomes.*
