## Supplemental Text for "On Copiotrophy and Temperature: Controls on Microbial Maximum Growth Rate Versus Translation Rate Optimization"

November 4, 2025

### S1 Text: Growth Model

#### S1.1 Text: Baseline Model

##### S1.1.1 Text: Baseline Model Introduction

As our starting point, we take the structure of the statistical model typically used to predict an organism's maximum growth rate from a genome sequence [5, 4]

$$\log(\mu_{\max}) \sim \beta_0 + \beta_1 T_{\text{opt}} + \beta_2 Q \tag{1}$$

where we let $Q$ be some genomic measure of translation optimization (usually the codon usage bias of highly-expressed genes like the ribosomal proteins; []). For practical prediction purposes $\mu_{\max}$ is often subjected to a Box-Cox transformation rather than a log transformation, but the use of the log transform will be useful below.

Since prediction is not our primary concern here, consider the analogous deterministic model

$$\mu_{\max}(T_{\text{opt}}, Q) = e^{\beta_0 + \beta_1 T_{\text{opt}} + \beta_2 Q} = e^{\beta_0} e^{\beta_1 T_{\text{opt}}} e^{\beta_2 Q}. \tag{2}$$

This model is very similar to the Arrhenius curve

$$\mu_{\max}(T_{\text{opt}}, Q) = e^{\beta_0} e^{\frac{\beta_1}{T_{\text{opt}}}} e^{\beta_2 Q} = A e^{\frac{-E_b}{RT}} \tag{3}$$

where $A = e^{\beta_0} e^{\beta_2 Q}$ is the pre-exponential factor and $\beta_1 = \frac{-E_b}{R}$, with the sole difference being that $\mu_{\max}$ is a function of $\frac{1}{T_{\text{opt}}}$ instead of $T_{\text{opt}}$. We will use the Arrhenius formulation in Eq 3 where $\mu_{\max}$ is a function of $\frac{1}{T_{\text{opt}}}$ for the remainder of our discussion, though our general results hold for Eq 2 as well. We assume temperatures are expressed in Kelvin.

For convenience, assume $Q$ is scaled such that $Q = 0$ indicates a complete lack of translation optimization and $Q \in [0, \infty)$. Then, our thermodynamic baseline expectation for maximum growth rate in the absence of translation optimization will be

$$\mu_0(T_{\text{opt}}) = e^{\beta_0 + \frac{\beta_1}{T_{\text{opt}}}}. \tag{4}$$

##### S1.1.2 Text: Inversely Related $Q$ and $T_{\text{opt}}$

Consider if $Q$ and $\frac{1}{T_{\text{opt}}}$ are linearly related, independently of any growth rate constraints, such that

$$\frac{1}{T_{\text{opt}}} = \beta_3 + \beta_4 Q. \tag{5}$$

Then,

$$\log(\mu_{\max}) = \beta_0 + \beta_1 (\beta_3 + \beta_4 Q) + \beta_2 Q \tag{6}$$

Assuming $\beta_1 < 0$ and $\beta_2 > 0$ (since we expect maximum growth rates to increase with both optimal temperature and translation optimization) and also that $\beta_4 > 0$ (such that optimal temperature and translation optimization are inversely related), we expect that the observed relationship between $\mu_{\max}$ and $Q$ will be negative when $|\beta_1 \beta_4| > \beta_2$.

##### S1.1.3 Text: Relative Fitness

We can write the relative fitness of a translation-optimized strain with maximum growth rate $\mu_{\max}$ as

$$\omega_Q = \frac{\mu_{\max}}{\mu_0} = e^{\beta_2 Q} \tag{7}$$

such that the log relative fitness is

$$\log(\omega_Q) = \log(\mu_{\max}) - \log(\mu_0) = \beta_2 Q. \tag{8}$$

Similarly, if instead we wish to calculate the relative fitness of a mutation conferring a change in translation optimization, $\Delta Q = Q_{\text{M}} - Q_{\text{WT}}$, where $Q_{\text{M}}$ is the translation optimization of the mutant and $Q_{\text{WT}}$ is the translation optimization of the wild-type, the relative fitness of that mutant can be expressed as

$$\log(\omega_{\Delta Q}) = \log(\mu_{\max}) - \log(\mu_0) = \beta_2 \Delta Q. \tag{9}$$

Thus, the relative fitness benefit of translation optimization does not depend on $T_{\text{opt}}$ following this model. Both psychrophilic and thermophilic organisms should derive the same degree of fitness benefit from a mutation conferring the same increase in translation optimization. Thus, this simple model by itself cannot explain the decreased evidence for genomic translation optimization among thermophiles (Fig 2, S2-4 Fig). It is also notable that a set amount of increase in $Q$ always leads to the same relative fitness of the mutant regardless of what the value of $Q$ was for the wild type following this model (*i.e.*, there are no diminishing returns for increasing $Q$).

Clearly, while this simple model formulation is sufficient for prediction tasks, it cannot explain more subtle patterns in trait variation among microbes such as the pattern of little genomic translation optimization in thermophiles relative to mesophiles (Figs. 2,3). In the next section we construct a model capable of capturing these patterns.

#### S1.2 Text: Modeling Diminishing Returns in Translation Optimization

##### S1.2.1 Text: Diminishing Returns Model

Consider a model where there is an exponential increase in an organism's maximum growth rate with increasing optimal growth temperature (as in S1.1 Text) and that organisms may additionally increase their maximum growth rate by increasing their translation optimization $Q$ (also as in S1.1 Text), but that the relative amount by which an increase in $Q$ increases $\mu_{\max}$ decreases with increasing $\mu_{\max}$ (diminishing returns). In other words, for fast growers, increasing translation optimization provides a smaller relative benefit than for slow growers. Such a model would be biologically reasonable given constraints on bacterial growth dynamics beyond translation rate limits. Even just considering translation itself, a variety of physical factors within the cell may constrain translation rates in addition to the degree of sequence optimization.

We express this saturation of the relationship between $Q$ and $\log(\mu_{\max})$ using a type II Holling response

$$\log(\mu_{\max}) = \beta_0 + \frac{\beta_1}{T_{\text{opt}}} + \frac{\beta_2 a \frac{Q}{T_{\text{opt}}}}{1 + a\frac{Q}{T_{\text{opt}}}} = \beta_0 + \frac{\beta_1}{T_{\text{opt}}} + \frac{\beta_2 a Q}{T_{\text{opt}} + aQ} \tag{10}$$

where $a$ is a scaling factor. In ecology, this type of curve is used to model the consumption rate of prey by a predator when there is some fixed handling time associated with each prey consumed [3, 2]. Here, we can similarly think about there being some handling time associated with translation that is invariant with $Q$ and $\frac{1}{T_{\text{opt}}}$. More complex saturating functions are possible (and may indeed better reflect the true relationship), but this simple formulation captures the general phenomenon of interest and is a convenient starting point.

As in S1.1 Text, we wish to calculate the relative fitness of a mutation conferring a change in translation optimization, $\Delta Q = Q_{\text{M}} - Q_{\text{WT}}$, where $Q_{\text{M}}$ is the translation optimization of the mutant and $Q_{\text{WT}}$ is the translation optimization of the wild-type. The log relative fitness of that mutant can be expressed as

$$\log(\omega_{\Delta Q}) = \log(\mu_{\max_{\text{M}}}) - \log(\mu_{\max_{\text{WT}}}) = \frac{a\beta_2 T_{\text{opt}} \Delta Q}{(T_{\text{opt}} + aQ_{\text{M}})(T_{\text{opt}} + aQ_{\text{WT}})}. \tag{11}$$

Consider the limits of this equation. As $Q_{\text{M}} \to \infty$ for fixed $Q_{\text{WT}}$ and fixed $T_{\text{opt}}$

$$\log(\omega_{\Delta Q}) \to \frac{\beta_2 T_{\text{opt}}}{T_{\text{opt}} + aQ_{\text{WT}}}, \tag{12}$$

as $Q_{\mathrm{WT}} \to \infty$ for fixed $\Delta Q$ and fixed $T_{\mathrm{opt}}$

$$\log(\omega_{\Delta Q}) \to 0, \tag{13}$$

and as $T_{\mathrm{opt}} \to \infty$ for fixed $Q_{\mathrm{M}}$ and fixed $Q_{\mathrm{WT}}$

$$\log(\omega_{\Delta Q}) \to 0. \tag{14}$$

Thus, (1) there is some maximum relative benefit to increasing $Q$ given a set starting value of $Q$ and fixed optimal growth temperature (Eq 10), (2) as the translation optimization of the wild-type strain increases to infinity the relative benefit of a mutation increasing $Q$ by a fixed amount decreases to zero (Eq 11), (3) and as optimal growth temperature goes to infinity the relative benefit of increasing $Q$ decreases to zero (Eq 12). In other words, the relative benefit of mutations increasing $Q$ decrease as the starting $Q$ and optimal growth temperature increase to infinity, *i.e.*, translation optimization will only get you so far.

Importantly, these relationships arise from a model that does not include any tradeoffs between growth rate and other traits (such as with substrate affinity). Such tradeoffs likely explain the great deal of variation among organism's value of $Q$ for a fixed $T_{\mathrm{opt}}$, especially for low $T_{\mathrm{opt}}$, but are not needed to explain the negative relationship between $Q$ and $T_{\mathrm{opt}}$.

##### S1.2.2 Text: Modeling A Hard Limit on Maximum Growth Rate

An alternative diminishing returns model posits some theoretical upper bound on maximum growth rates, $\mu_U$, above which no organism can go. Let

$$\mu_0[T_{\mathrm{opt}}] = e^{\beta_0 + \frac{\beta_1}{T_{\mathrm{opt}}}} \tag{15}$$

be the baseline maximum growth rate when $Q = 0$.

We can then write our upper-bound model as

$$\mu_{\max} = \mu_0[T_{\mathrm{opt}}] e^{\log\left(\frac{\mu_U}{\mu_0[T_{\mathrm{opt}}]}\right)\left(\frac{aQ}{1+aQ}\right)} \tag{16}$$

assuming $Q \in [0, \infty)$ and $\mu_0[T_{\mathrm{opt}}] \leq \mu_U$ (for example, if we assume an upper bound $T_U$ on $T_{\mathrm{opt}}$ such that $\mu_0[T_U] = \mu_U$). Under this model, $\mu_{\max} \in [\mu_0[T_{\mathrm{opt}}], \mu_U]$ and higher values of Q increase $\mu_{\max}$. This model has the same number of parameters as the model in S1.2.1 Text, since we have added one parameter ($\mu_U$) and removed one parameter ($\beta_2$).

Similar to above, we can calculate the relative fitness of a mutant strain as

$$\log(\omega_{\Delta Q}) = \log\left(\frac{\mu_U}{\mu_0[T_{\mathrm{opt}}]}\right)\left(\frac{a\Delta Q}{(1 + aQ_{\mathrm{M}})(1 + aQ_{\mathrm{WT}})}\right) \tag{17}$$

with similar overall model behavior.

#### S1.3 Text: Selection Coefficients

##### S1.3.1 Text: Selection Coefficients for Non-Saturating $Q$

We can use the relative fitness to calculate the selection coefficient for a particular degree of $Q$ relative to the thermodynamic baseline

$$s_Q = \omega - 1 = e^{\beta_2 Q} - 1. \tag{18}$$

Chevin [1] notes that calculating selection coefficient in this way, while common among microbiologists (especially in the field of experimental evolution), is not technically correct. Instead, Chevin suggests that one calculate the per-generation selection coefficient for organisms replicating via binary fission as

$$s_Q = \left(\frac{\mu_{\max} - \mu_0}{\mu_0}\right)\log(2) = \left(e^{\beta_2 Q} - 1\right)\log(2). \tag{19}$$

Similarly, one can calculate the selection coefficient for a mutation conferring a change in translation optimization $\Delta Q = Q_{\mathrm{Mut}} - Q_{\mathrm{WT}}$ as

$$s_{\Delta Q} = \left(e^{\beta_2 \Delta Q} - 1\right)\log(2). \tag{20}$$

##### S1.4.2 Text: Selection Coefficients for Saturating $Q$

As in S1.3.1 Text, one can calculate the selection coefficient of a mutation conferring a change $\Delta Q$ from a wild-type strain with translation optimization $Q_{\mathrm{WT}}$ as

$$s_{\Delta Q} = \left(e^{\frac{a\beta_2 T_{\mathrm{opt}}\Delta Q}{(T_{\mathrm{opt}}+aQ_{\mathrm{M}})(T_{\mathrm{opt}}+aQ_{\mathrm{WT}})}} - 1\right)\log(2). \tag{21}$$

[5] J. L. Weissman, Shengwei Hou, and Jed A. Fuhrman. Estimating maximal microbial growth rates from cultures, metagenomes, and single cells via

codon usage patterns. *Proceedings of the National Academy of Sciences*, 118(12):e2016810118, March 2021. Publisher: Proceedings of the National Academy of Sciences.
